# Functional Asymmetries between Central and Peripheral Retinal Ganglion Cells in a Diurnal Rodent

**DOI:** 10.1101/277814

**Authors:** María-José Escobar, Mónica Otero, César Reyes, Rubén Herzog, Joaquin Araya, Cristóbal Ibaceta, Adrián G. Palacios

## Abstract

The segregated properties of the visual system processing central or peripheral regions of the visual field have been widely studied in the visual cortex and the LGN, but rarely reported in retina. The retina performs complex computational strategies to extract spatial-temporal features that are in coherence with animal behavior and survival. Even if a big effort has been done to functionally characterize different retinal ganglion cell (RGC) types, a clear account of the particular functionality of central and peripheral cells is still missing. Here, using electrophysiological data obtained with a 256-MEA recording system on female diurnal rodent retinas (*Octodon degus*), we evidenced that peripheral RGCs have larger receptive fields, more sustained, faster and shorter temporal responses and sensitive to higher temporal frequencies with a broader frequency bandwidth than the center. Additionally, we also compared the asymmetries between ON and OFF cell populations present in each region, reporting that these asymmetries are dependent on the eccentricity. Finally, the presence of the asymmetries here reported emphasizes even more the complexity of computational strategies performed by the retina, which could serve as inspiration for the development of artificial visual systems.

## 1. Introduction

The segregated properties of the visual system processing central or peripheral regions of the visual field have been widely studied in the visual cortex and the LGN (Stone, 1983; Orban et al., 1986; Loschky et al., 2017), but rarely reported in the retina.

In general, the distribution of cells is not homogeneous along the retina and it varies with the eccentricity. In rodents, retinal ganglion cells (RGC) are concentrated at the center of the retina forming a visual stripe, see mouse (Dräger and Olsen, 1981); albino and pigmented mice (Salinas-Navarro et al., 2009a); and rats (Salinas-Navarro et al., 2009b). In pigmented rabbits the RGC soma diameter varies with the eccentricity (Provis, 1979) and the retina of the archerfish revealed a spatial organization locating receptive fields (RF) with high resolution at the central region (Ben-Simon et al., 2012). An interesting case is the diurnal rodent *Octodon degu* (Chávez et al., 2003), who showed isodensity zones where the highest concentration of RGCs is located at the central retina (Vega-Zuniga et al., 2013), what seems to match the spatial distribution of retina photoreceptors (Jacobs et al., 2003).

The functional characterization of RGCs, obtained from the analysis of cell response parameters such as latency (short / long), ON / OFF, sustained / transient, biphasity or receptive field properties has revealed a huge diversity (Carcieri et al., 2003; Heine and Passaglia, 2011; Baden et al., 2016) indicating a large number of functional channels originated at the RGC level. This functional variety of RGCs could be explained observing the circuitries and anatomical cell types identified by morphological, immunomarker and molecular criteria (Euler et al., 2014; Masland, 2012b).

Interestingly, functional data in non-human primates proposes regular mosaics of a single functional cell type, paving the entire visual field (Gauthier et al., 2009; Shlens et al., 2009). Nevertheless, this result has not been validated in other animal species using a functional type characterization based on linear non-linear models. With a functional characterization based in information theory (Sharpee, 2013, 2014), Kastner and Baccus (2011) reported functional mosaics in the salamander retina. Even if some of the RGC anatomical types seems also to pave the entire retinal mosaic, their functional characterization can vary according to several factors including retinal eccentricity, life style, micro-habitat; behavioral functionality. A recent study in primate retina emphasizes the differences in the electrophysiological response of a single RGC anatomical type (Midget) located in foveal, center and peripheral retina regions (Sinha et al., 2017; Masland, 2017), revealing different functional properties associated to a single RGC type. However, different functional properties for the entire RGC populations remain unknown together with the asymmetries present between ON and OFF cell population at different locations of the visual field (Chichilnisky and Kalmar, 2002; Zaghloul et al., 2003; Liang and Freed, 2010; Jiang et al., 2015; Leonhardt et al., 2016).

Here we report for in a diurnal rodent *(Octodon degu*) the functional asymmetries between central and peripheral RGCs in terms of the number of cell types; RF sizes, temporal response kinematics and temporal frequency properties. We detect important functional asymmetries between the two regions, suggesting different ecological meanings and projections to higher visual areas. Additionally, we also evaluated the asymmetries between ON and OFF cell populations for each retina region dissected. Functional asymmetries are present between these two cell populations, and more importantly, these asymmetries depend on the eccentricity being hard to propose a general rule. We also evaluated a pair of representative RGCs of the central and peripheral retina as image filters, observing the variety of information associated to each of them, presenting the retina computation strategies as a source of inspiration for the design of artificial vision devices.

## 2. Materials and Methods

### 2.1. Multielectrode recording and animal preparation

Adult young female, 3-8 months old, *Octodon degus* born in captivity and maintained in large collective cages in an adapted animal facility at 2025^*°*^ C, 12-h / 12-h light-dark cycle, with access to pellet food and water *ad-libitum*. Experiments were carried after approval by a bioethic committee at the Universidad de Valparaiso in accordance with regulation of the Chilean Research Council (CONICYT).

The experimental recording methodology was already described in Palacios-Muñoz et al. (2014). In brief, for extracellular multi-electrodes recording array (MEA-256-USB Multichannelsystem, Germany) experiments, the animals were dark-adapted for 30 minutes and deeply anesthetized with Isoflurane (Sigma-Aldrich Co.) previous to euthanizing by decapitation. Eyes were quickly enucleated and eye-cup prepared under dim red light and immersed in bicarbonate-buffered Ames Medium (Sigma-Aldrich Co.), kept constantly oxygenating (95% + 5%) at room temperature. Retina pieces, following the special orientation of optic nerve (it is slipping to dorsal and nasal zone), getting a central and periphery zones (dorsal and ventral). The dissected center part of the retina was obtained cutting a spot in central zone next to optic nerve in the temporal side. On the other hand, a peripheral piece of the retina was obtained from middle of the dorsal zone (Fig 1A).

**Figure 1:**
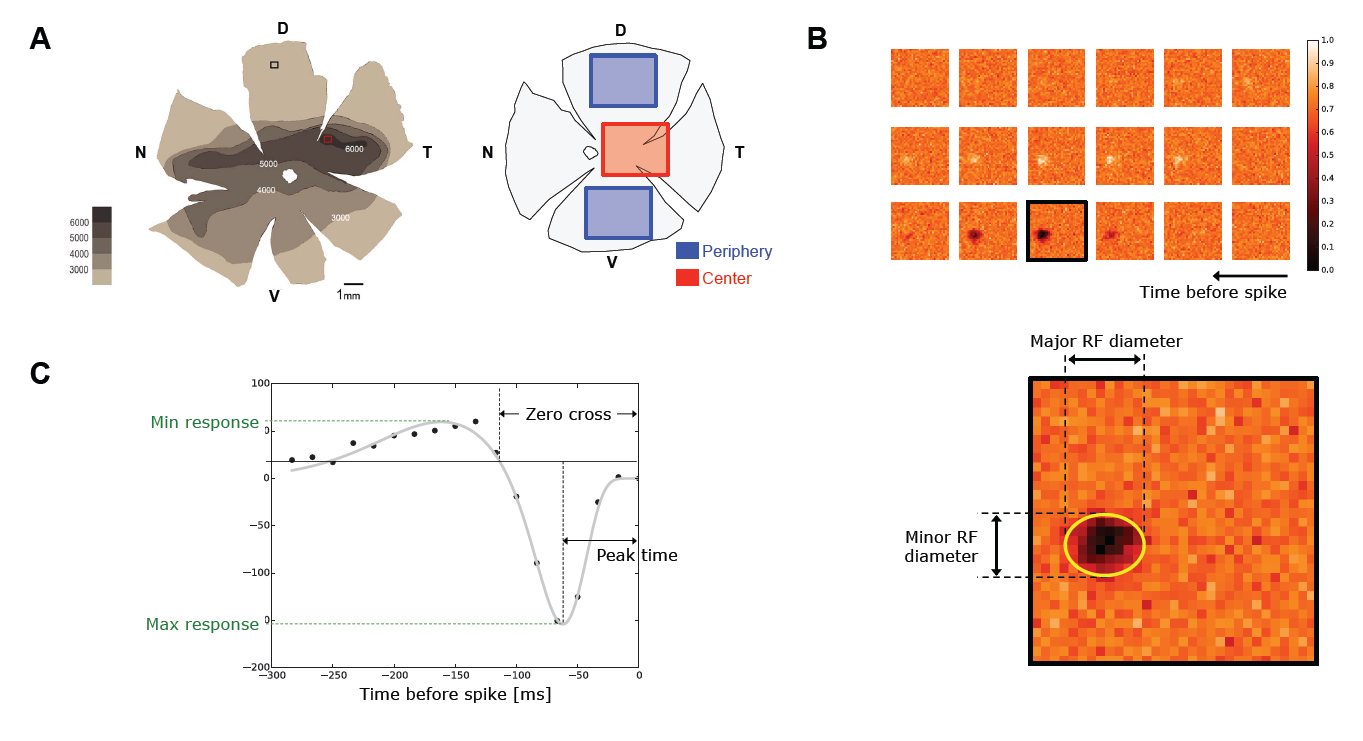
**A** (Left) *Octodon degu’s* retinal wholemount showing the ganglion cell layer density map for the left eye retina reported by Vega-Zuniga et al. (2013)). (Right) Retina diagram specifying the dissected regions considered as central and peripheral retina (red and blue boxes, respectively). **B** Sample of the time course STA obtained for an OFF RGC. Parameters used for the spatial characterization, such as minor and mayor RF diameters, are obtained fitting an ellipse on the frame with the highest activity. **C** Temporal time course of the same OFF cell shown in **B**. Each black dot represents the intensity around the center of the RF for each frame. Gray line shows the resulting fitting curve (see equation (1)). From the fitted version we extracted parameters such as, peak-time, zero-cross and maximal and minimal response.

Small pieces of the retina were left aside in darkness during others 15min to stabilize, them were carefully separated from retinal pigment epithelium and mounted on a ring containing a dialysis membrane (Spectra / Por 6, MWCO 25,000) which was fit in an up / down plastic cylinder that was adjusted to contact the retina (RGCs layer) with the MEA surface. Multi-electrodes array (MEA USB-256, Multichannel Systems GmbH, Germany) for in vitro isolated retinae were used to record compound action potential from RGCs under dark adapted conditions

### 2.2 Visual stimulation

Visual stimuli were generated in a customized software created in PsychoToolbox (MatLab) in a Mac mini desktop computer (Apple) and were projected onto the retina by a LED projector (PLED-W500, ViewSonic, USA) equipped with an electronic shutter (Vincent Associates, Rochester, USA), trough an optical system, with a series of optical density filters to control light intensity, connected to an inverted microscope (Lens 4, Eclipse TE2000, NIKON, Japan) and a CCD camera (Pixelfly, PCO, USA) for visualization and calibration for each stimulus projection.

Light flashes produced with full screen stimulation with a period of 1400ms divided in 400ms ON and 1000ms OFF. The light flash sequence was repeated 30 times. The projected images were conformed by 372×372 pixels, where each pixel covers a surface of 16um^2^. Since rodents are dichromatic (green and blue / UV cones), only B (blue) and G (green) beams of the projector were used as visual stimulation, while the R (red) channel was used for signal synchronization. Checkerboards were formed by black combined with B and G colors with a bin size of 50um at a rate of 60fps for 30min. All parameters of the stimuli were programed by a custom software in Matlab.

### 2.3. Spike sorting and data analysis

Action potentials were recorded using MEA trough the MC / Rack software (MultichannelSystem, Germany) and data analyzed by the SpykingCircus software (Yger et al., 2016). This software help a simultaneous spike sorting from hundreds of neurons, which are checked using a dedicated GUI to remove duplicities and those units, which did not have the shape of a spike. Each unit was manually checked and confronted with similar entities with the same activity pattern to avoid duplicities. The spikes obtained were validated using inter spike interval (ISI) and the autocorrelation general criteria. Also, it was checked the cross-correlation between previously obtained templates. Those templates with a high cross correlation and similar receptive field characteristics and light responses were merged.

Regularly 200-300 RGCs were isolated from a single experiment. RGCs responses were then analyzed using custom software spike sorting and then for receptive field estimation.

### 2.4. Receptive field estimation

For all the stimuli, here analyzed we collected data from four retinal pieces (4 different animals) conforming two central and two peripheral regions to a total of 659 RGCs: 390 for center and 269 for periphery, respectively.

For estimating a receptive field (RF) a checkerboard stimulus wae used. The spatio-temporal receptive field (RF) of each cell was estimated using custom-built software, which performs Spike Triggered Average (STA) (Chichilnisky and Kalmar, 2002). STA was computed for each of the 659 RGCs recorded, the spatial and temporal profile of the estimated linear RF was obtained.

The spatial component of the RF was obtained fitting a 2D Gaussian on the frame with maximal response. The temporal component of the RFs was recovered collecting the average response of a window of 3×3 pixels around the center of the spatial RF along 18 frames prior the action potential generation. The temporal profile was then fitted by a difference of two cascade low-pass filters as it is shown in Chichilnisky and Kalmar (2002).

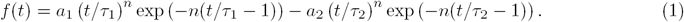

### 2.5. Light response characterization

RGCs were analyzed according to their response to light flashes stimulation. The RGC firing rates was used to classify the neurons in ON, ON-OFF or OFF cells. To do so, we calculated the Peristimulus Time Histogram (PSTH) with a bin size of 5ms, to each cell in order to determine the relationship between timing of the ON and OFF flashes presentation inter trials. For each PSTH we computed the mean (*μ*) and the standard deviation (*σ*) response along the stimulus period. Only units with a significant activity over *δ* = *μ ±* 1.5*σ* along the ON and OFF flashes were selected, and the threshold *δ* was also used to discard those cells with a very low activity (less than 20 spikes in all bins).

The latency of each cell response was computed as the time when the maximal firing rate is reached after the stimulus onset. The sustained index used to analyze different light-flashes responses were computed as

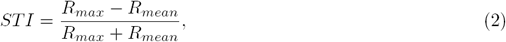

where *R*_*max*_ (*R*_*mean*_) is the maximal (mean) firing rates of the cell observed in a given condition.

To obtain the different groups of cell types according to their temporal response, we used the fitted curve defined in (1). The fitted curve allowed us to extend the temporal characterization of the cell up to 40 frames prior the spike generation (666ms) with a resolution of 900 points. Starting from this generated curve, we calculate some temporal characteristics of the RF such as: peak time, peak amplitude, zero crossing, biphasic index and roundness. The peak time represents the moment at the maximal contrast intensity. The zero-crossing of the curve stands for the difference between the generation of the spike and the zero-crossing time of the curve. The peak amplitude is only the value of the STA contrast at the peak time. Moreover, the biphasic index is the absolute value of the ratio between the minimal and maximal amplitude of the STA time curve (see Fig. 1**C**). In the other hand, the roundness is related to the ratio between the minor and major semi-axes of the RF ellipse.

Temporal cell profiles were clustered using the five first principal components (PCA) decomposition. Following the approach proposed by Rodriguez and Laio (2014), which sets as centroids of the clusters the templates with the highest density, we obtained a total of 17 clusters for central and 14 clusters for peripheral retina. Once each cluster has been obtained, we computed the mean of the temporal profiles for each of them obtaining also the parameters of the fitted curve defined in (1). As the Fourier transform (FT) of (1) is defined, the fitted parameters were also used to obtain a temporal frequency domain representation of each temporal cluster.

### 2.6. Linear and Non-linear response

For each cell, linear response was computed convolving the estimated RF (obtained with STA) with the checkerboard stimulus. The curve *n*_*l*_ relating linear with real response was fitted by the following curve

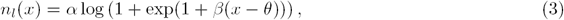

where mainly the parameters *β* and *θ* quantify the nonlinearity represented in the sigmoid function described in (3).

## 3. Results

To characterize different cell populations of RGCs present in the *Octodon degus* retina, we collected a series of data coming from *ex-vitro* recordings in a 256-multi electrodes array system. We recorded the RGC response confronted to two types of visual stimuli: light flashes and white-noise checkerboards.

We analyzed four pieces of different retinas dissecting regions following RGC isodensity distribution, as it was reported by Vega-Zuniga et al. (2013) (see Fig. 1**A** left). We dissected three central areas and two peripheries (see Section Methods) counting a total of 809 RGCs (389 / 264 for two center / periphery, respectively). Only cells with valid receptive fields (STA) were considered in the analysis. Here we show spatial and temporal characteristics of the two retina regions, observing significant difference between them.

### 3.1. ON, OFF and ON-OFF cell populations

Light flashes of 400ms were used to establish ON / OFF characteristic of RGCs. A PSTHs plot was binned in time windows of 5ms length. Cells with peak responses higher than the mean response, plus 1.5 times the standard deviation, were classified as ON, OFF or ON-OFF populations. 1 shows detailed values of the distribution percentage of each cell type, for each retina piece.

**Table 1:**
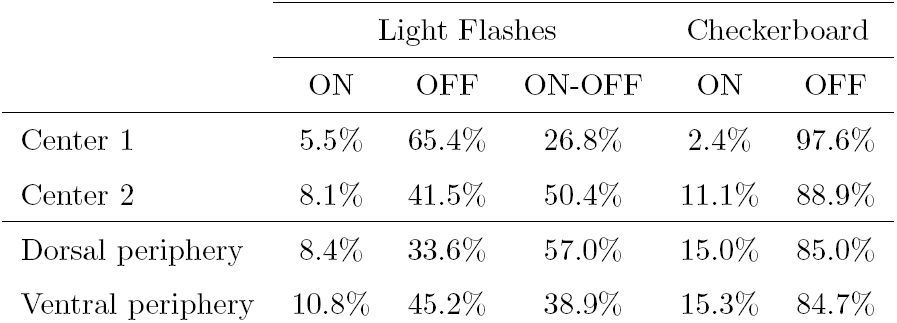
Percentages of ON, OFF and ON-OFF cells for central and peripheral retina.

|  | Light Flashes |  |  | Checkerboard |  |
| --- | --- | --- | --- | --- | --- |
|  | ON | OFF | ON-OFF | ON | OFF |
| Center 1 | 5.5% | 65.4% | 26.8% | 2.4% | 97.6% |
| Center 2 | 8.1% | 41.5% | 50.4% | 11.1% | 88.9% |
| Dorsal periphery | 8.4% | 33.6% | 57.0% | 15.0% | 85.0% |
| Ventral periphery | 10.8% | 45.2% | 38.9% | 15.3% | 84.7% |

Additionally, using white noise checkerboard as input stimulus we computed the spike-triggered average (STA), for each RGC, obtaining the temporal profile and spatial extent of their responses. RGCs were categorized in ON and OFF response populations according to the polarity of their time course. We observe no significant difference in the proportion of ON and OFF cells between central and peripheral retina (see 1), which values are similar as the ones reported in (Palacios-Muñoz et al., 2014)

### 3.2. Sustained versus transient response

We additionally measured the temporal course response of each RGC confronted to light flash stimulus. According to cell response to light flashes they were classified in ON-OFF, ON or OFF, with a temporal response profile as shown in Fig. 2-left column (refernce image obtained from dorsal periphery data). For the ON, OFF and ON-OFF cell populations we computed the latencies and the sustained index (see Methods, (2)). Fig. 2**A** quantifies the latencies values obtained for the four pieces of retina analyzed, separated by response type. Response latencies of central RGCs are represented in red while peripheral values in blue. In the case of ON-OFF cell population (top) we observed a variety of responses most of them clusterized and probably attributed to a single cell type. Central RGCs of ON-OFF cell population present larger latencies for the OFF response compared to peripheral values (top, p*<*1e-4 Kolmogorov-Smirnof), which is also observed in the OFF populations (bottom, p*<*1e-8 Kolmogorov-Smirnof). However, the ON response has divided behaviors: only the ON-OFF cell population (top) presents differences between central and peripheral cells (p*<*0.01, Kolmogorov-Smirnof), while ON cell population (middle) only present significant differences for one of the centers analyzed (Center 1, p*<*1e-8, Kolmogorov-Smirnof), but not for the other (Center 2, p*>*0.1, Kolmogorov-Smirnof).

**Figure 2:**
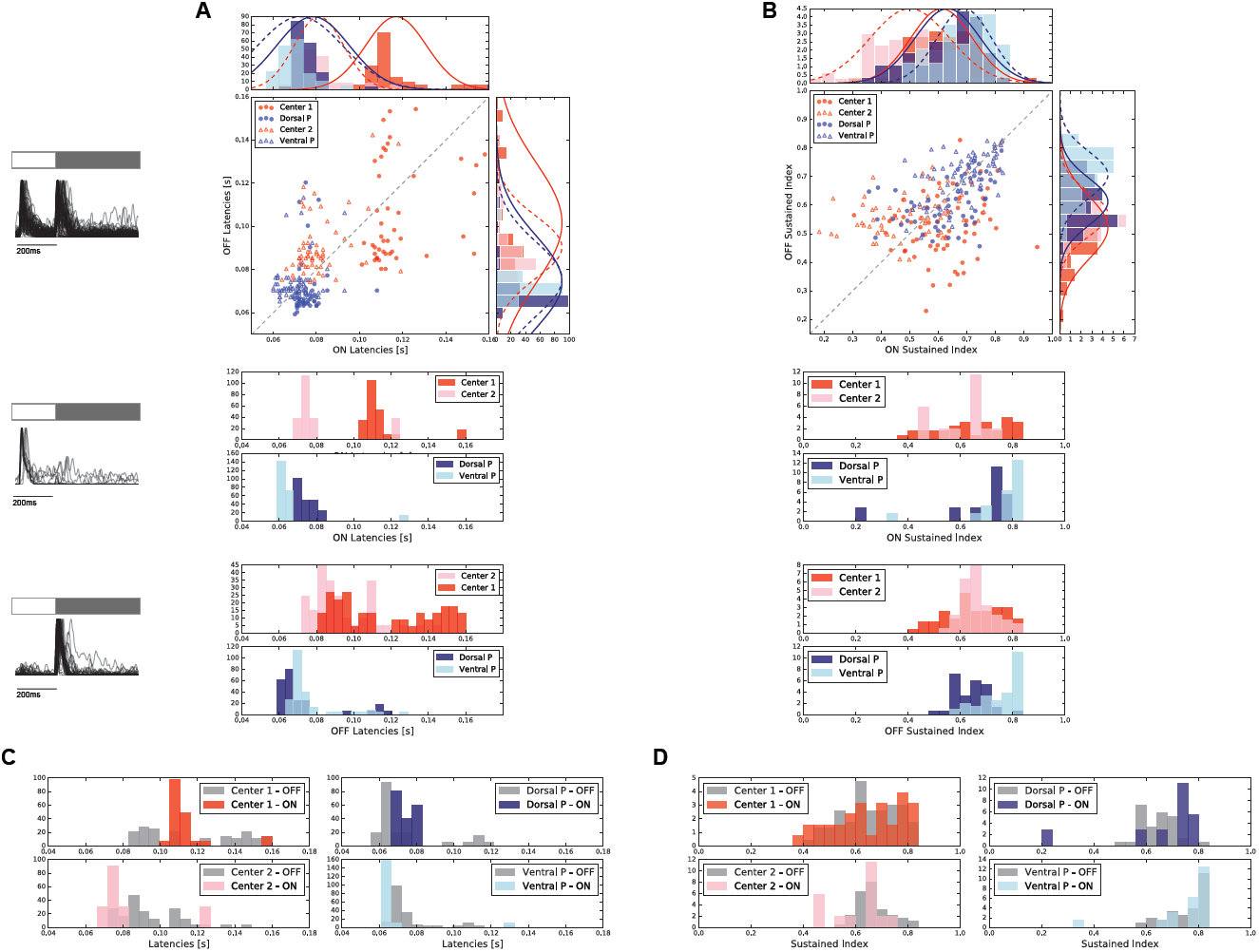
Different flashes responses obtained for central and peripheral retina. A condensed representation of the PSTHs of all the RGCs associated to each cell population, for the dorsal peripheral retina in shown in the left. **A**) Response latencies observed in each cell population for the four retina pieces analyzed, for the ON-OFF (top), ON (middle) and OFF (bottom) cell populations. **B**) Sustained index observed in each cell population for the four retina pieces analyzed, for the ON-OFF (top), ON (middle) and OFF (bottom) cell populations. **C**) Comparison of response latencies between ON and OFF cell populations in the four pieces of retina analyzed. **D**) Comparison of sustained index (SI) between ON and OFF cell populations in the four pieces of retina analyzed.

Cell responses were also compared in terms of their sustained or transient response temporal courses (see Fig. 2**B**). Sustained cell responses have a large sustained index (SI) value, which is the contrary for transient temporal cell responses. In terms of differences between center and periphery, in ON-OFF cell population (Fig. 2**B**-top) the OFF peak has a higher sustained response compared with the center (Kolmogorov-Smirnof p*<*0.01), which is not always held for the ON peak (center 1 with Dorsal P, Kolmogorov-Smirnof p*>*0.4; p*<*0.02 for all the other cases). By separated, ON and OFF cell populations (Fig. 2**B**-middle and bottom, respectively), showed similar behavior. ON cell population exhibited a more sustained response in the periphery compared to central retina (Kolmorov-Smirnof, p*<*0.01; with the exception of Center 1-Dorsal P, p*>*0.3), which is also held for OFF cell population.

Additionally, we evaluated asymmetries between ON and OFF cell populations and how these asymmetries vary depending on retina eccentricity. The 2D charts presented in Fig. 2**A** and **B**, shows for a single retina piece, the comparison between the ON and OFF peak response of the ON-OFF cell populations. We observe a variety of results. For instance, Center 1 has ON peaks with a larger response latency and SI (p*<*1e-5) compared to OFF response, while Center 2 presents the opposite effect where the OFF peaks presents the largest response latency and SI (p*<*1e-5). Similarly, the results found in the peripheries is also antagonist: dorsal periphery has ON response with a larger response latency compared to OFF response, and ventral periphery reports the opposite. Nevertheless, no differences where observer between the SI of ON and OFF responses in the ON-OFF cell population. We now contrasted response latencies and SI for the ON and OFF cell populations by separated. Fig. 2**C** shows the comparison of latencies for the four pieces of retina analyzed. Similarly to ON-OFF cell population, center 1 presents shorter latencies for OFF response (p*<*0.004) while center 2 evidences the opposite (p*<*0.001). Same case for the peripheries: dorsal periphery has faster OFF cells (p*<*0.001) and ventral periphery faster ON cells (p*<*1e-5). Fig. 2**D** shows similar analysis by now concerning the SI, where now only the dorsal periphery showed significant differences between ON and OFF cell populations (p*<*0.01).

### 3.3. Spatial functional properties

The RGC response to white-noise stimulus was used to compute the STA of each cell. We only considered cells with a valid STA in the analysis. The spatial component of the STA RF was fitted with a 2D Gaussian to the frame representing the temporal peak-response of the cell, as it is specified in the Methods and shown in Fig. 1**B**.

We analyzed the spatial properties of the central and peripheral retina regions dissected, as shown in Fig. 3. For each dissected piece we computed the map of all the RGC RF encountered (sample maps Fig. 3**A**). As the center region has strong variability in the RGC density (Vega-Zuniga et al., 2013; Jacobs et al., 2003), we computed the variability of the RGC sizes along the dissected piece to detect if we are placed over a homogeneous zone. The graphs located at the top and left of the cell maps, show the mean RF area (red and blue dots, respectively) along all the RGCs located at a certain *x* or *y* position of the array, respectively. Additionally, gray bars represent the number of RGCs found in that position which are labeled in the right axis of the graph. As we can observe, center region seems homogeneous (x-axis: r-value = 0.31, p*<*0.15; y-axis: r-value =-0.09, p*<*0.7) in contrast to the dorsal peripheral region who presents a variation of the mean RF area along the y-axis (x-axis: r-value = 0.06, p*<*0.8; y-axis: r-value = 0.56, p*<*0.01).

**Figure 3:**
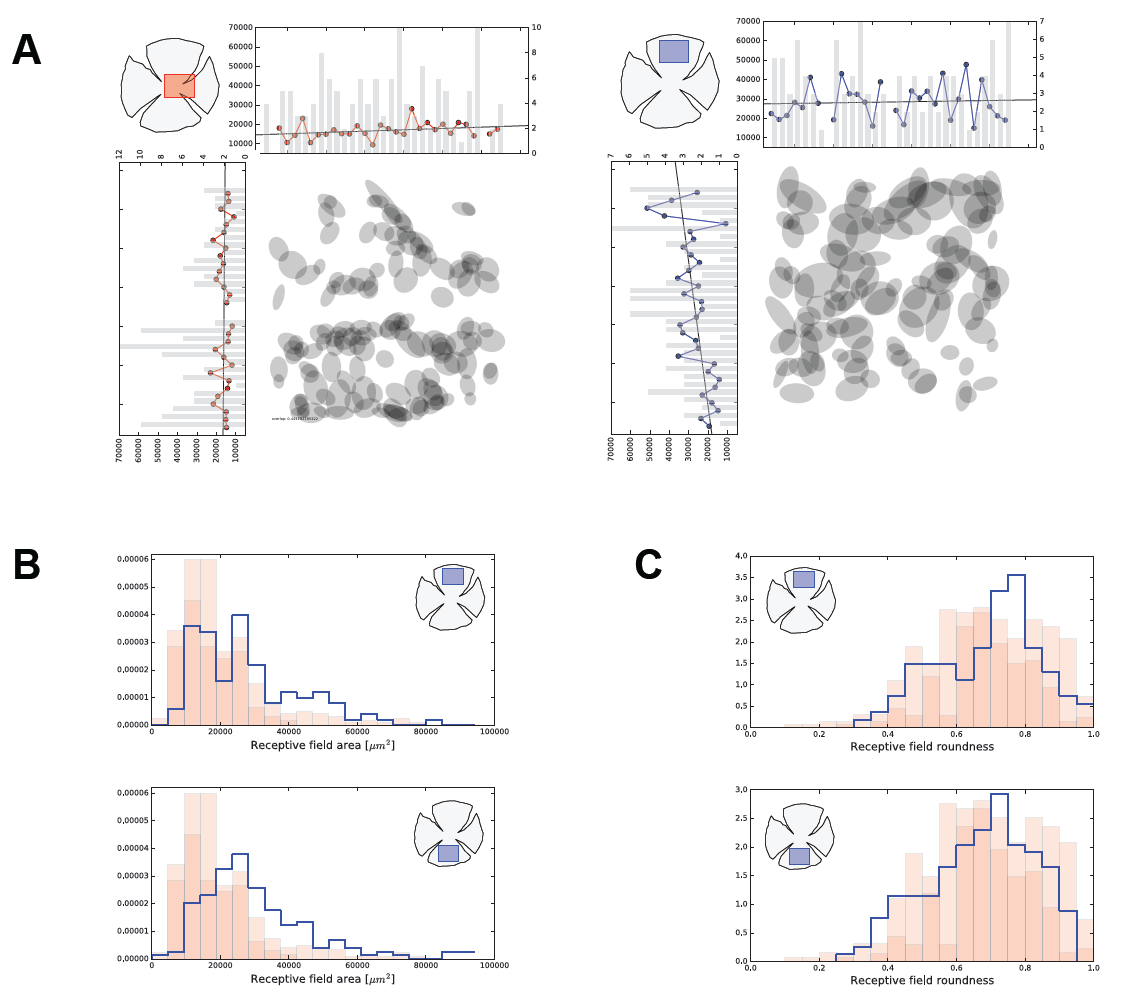
Spatial features observed in central RGCs. **A**) Maps of RGC RF encountered in one sample of central (left-red) and peripheral (right-blue) retina studied. Each map is accompanied with two charts indicating the variation of RF areas along the *x* and *y* axis, emphasizing the fact that central retina areas contain a high variability in RGC number. **B**) Distribution of RF areas of the central (red bars) versus peripheral retina (stepped blue histogram). **C**) Distribution of RF roundness of the central (red bars) versus peripheral retina (stepped blue histogram).

In general, RGCs haver larger RF for peripheral areas compared to the central areas of the retina (see Fig. 3**B**). We compared the distribution of RF area for the different tissues studied. Each histogram shows the distribution of the RF area encountered for each retina tissue (red for center and blue for periphery). The two centers have smaller RF areas compared to dorsal and ventral peripheries (Kolmogorov-Smirnof, p¡0.001 and p¡1e-8, respectively).

Most of the cells have elongated RF, as can be observed measuring the RF roundness parameter (Fig. 3**C**). We computed the RF roundness for each cell dividing the minor versus the major RF diameter (see Fig. 1). Similarly, no statistical differences were observed between ON and OFF cell populations. In addition, we also compared the distribution of roundness between center and periphery obtaining significant differences with the centers only for ventral retina (Kolmogorov-Smirnof p*<*0.01 and p*<*0.005).

### 3.4. Temporal functional properties

Temporal analysis obtained from STA reflected different cell dynamics in central and peripheral retina. Each RGC temporal course profile was obtained as the intensity value of the RF center along a time window of 400ms previous the spike emission. The temporal profile was fit with a mathematical function (see Section Methods, Chichilnisky and Kalmar (2002)), providing a parametric expression where the parameters of interest can be analytically extracted.

Both, for center and periphery we clusterized the temporal profiles, by experiment, obtaining the representation shown in Fig. 4**A**. The color of the temporal profile indicates the associated cluster, which average value is represented with a thick colored line. We show as example only one center with 7 clusters (left), and a ventral periphery with 15 cluster (right). From the parametric expression obtained for each RGC time course, we extracted values of interest such as peak-time, zero-cross and biphasic index (see Fig. 1**C**).

**Figure 4:**
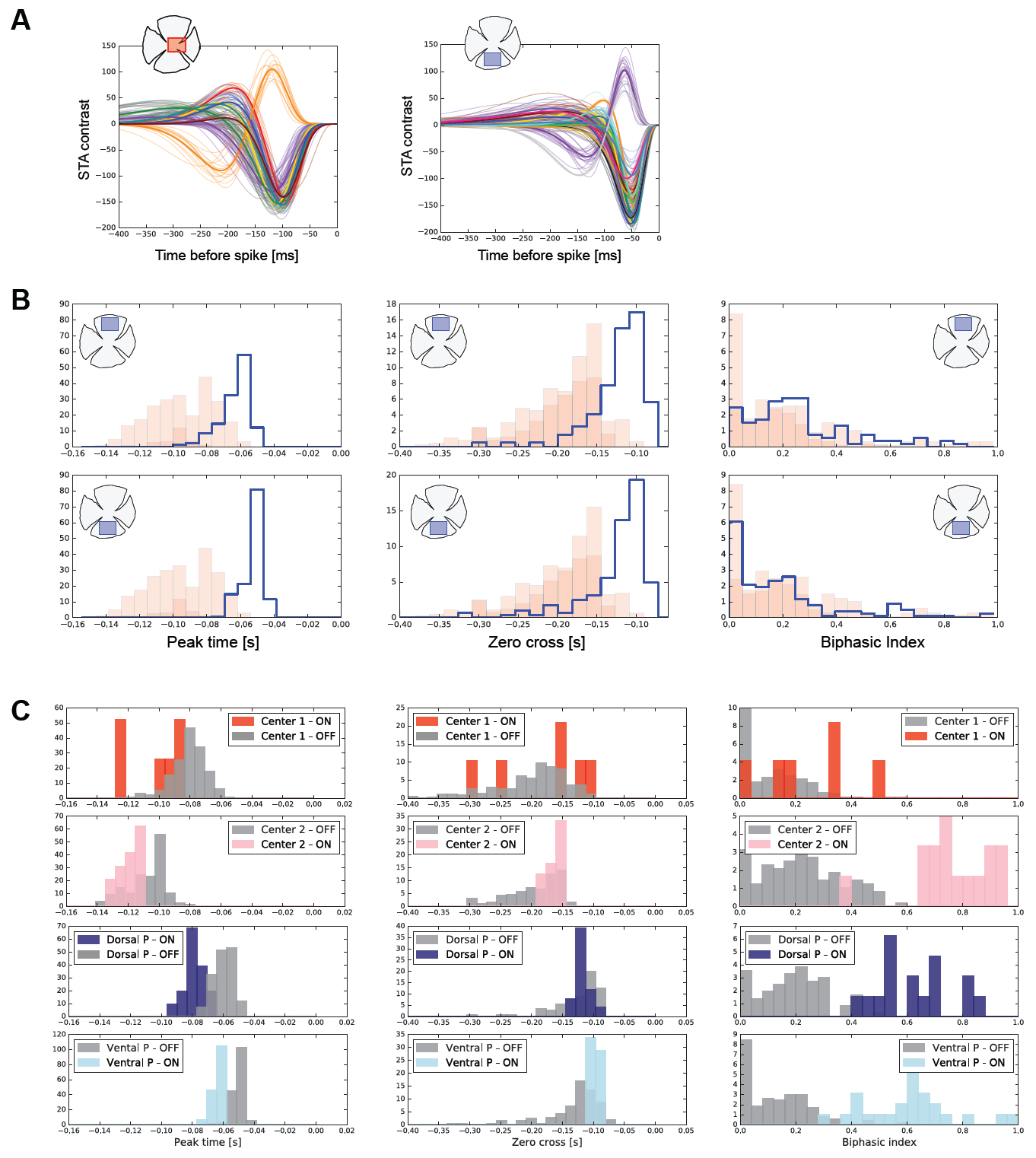
Temporal features observed in central RGCs. **A**) Temporal profile of all the RGCs placed at one of the center studied (left) and one periphery (right). Each color line represents the associated cluster, which mean curve is plotted with a thick colored line. The number of clusters encounters, from top to bottom, 7 and 15, respectively. **B**) Distributions of the peak-time (left), zero-cross (middle) and biphasic-index (right) parameters for the total number of retina pieces analyzed (two centers and two peripheries). **C**) Distributions of the peak-time (left), zero-cross (middle) and biphasic-index (right) parameters for the total number of retina pieces analyzed contrasting ON versus OFF cell populations.

Fig. 4**B** shows the comparison of temporal properties of central versus peripheral RGCs. For the peak-time (left), both peripheries are faster than any of the central regions studied (Kolmogorov-Smirnof, p¡1e-5), the duration of the temporal response represented by the zero-cross (middle) parameter is also shorter for RGCs located at the periphery (Kolmogorov-Smirnof, p*<*1e-5). In the case of biphasic index (right), only one pair of center-periphery (dorsal-center 2) showed no significant differences (Kolmogorov-Smirnof, p*>*0.01).

Not only differences between central and peripheral RGCs temporal properties are observed, but also, between ON and OFF cell populations. Fig. 4**C** shows a comparison between temporal properties of ON and OFF cell populations belonging to the same retina tissue. In all the experiments we observe than OFF cell population has lower peak-time values compared to ON cells. A similar behavior is observed in the zero-cross parameter. Both centers exhibit lower zero-cross values for ON cell population. A fast ON cell population is also observed in the ventral periphery, while no significant differences were observed between ON and OFF cell population in the dorsal retina (p*>*0.1). Regarding biphasic index, in all the tissued analyzed, we observed a notable difference between both cell populations: ON cells have a more biphasic response compared to OFF cells.

### 3.5 Frequency selectivity of central and peripheral retina

Frequency response of each RGC was obtained from the Fourier transform of the fitted RF temporal responses described in (1). Fig. 5**A** shows the magnitude of the Fourier transform of each RGC, normalized to the same peak amplitude, for centers and peripheries. We firstly focused on correlate time domain feature parameters with temporal frequency characterization. Fig. 5**B** represents the relation between the zero-cross value of the temporal response with frequency selectivity. Low-pass filter RGCs (cells with frequency selectivity at 0[Hz]) are observed in all over the range of possible zero-cross values. The green area of interest is zoomed on the right emphasizing the differences encountered between RGCs in central (red) and peripheral (blue) regions showing a correlation between these two values (r-value = 0.73, p-value*<*1e-5). Basically, cells at the central retina have larger temporal responses and lower frequency selectivities compared to the periphery. In addition, cells with the same zero-cross values have higher frequency selectivities in the center compared to peripheral region. Interestingly, ON cell population both in central and peripheral retina have the highest frequency selectivities (see Fig. 5**B** right), and none of them acts as a low pass filter with zero frequency selectivity.

**Figure 5:**
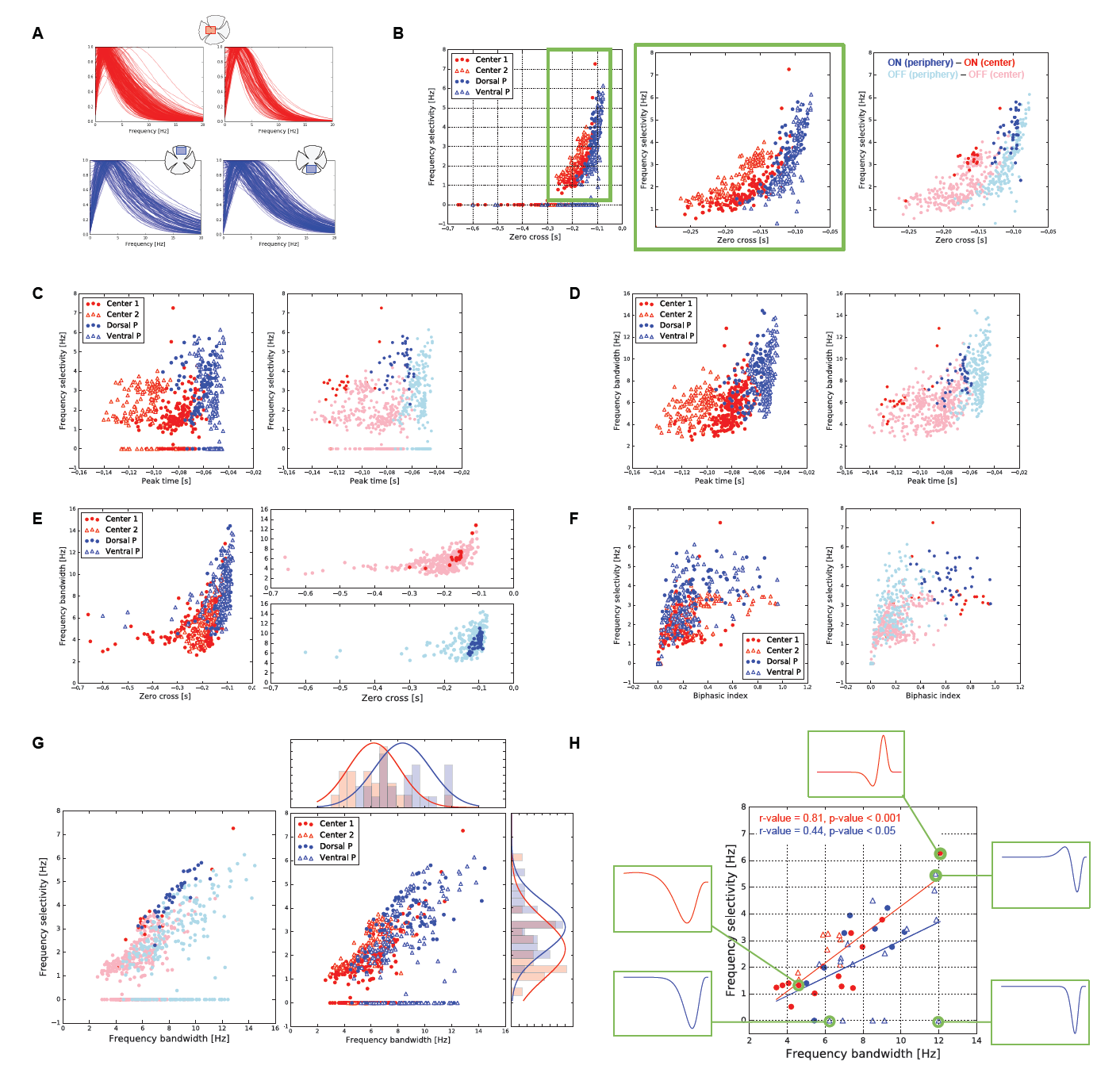
Frequency selectivity in central and peripheral RGCs. **A**) Frequency response of the RGCs temporal profiles for cells placed at the retina center (red) and periphery (blue). **B**) Relation between temporal zero-cross parameter and frequency selectivity. The green box marked on the left graph is zoomed on the middle graph. We also show how these two parameters are related to ON and OFF cell populations (right). Both, center and periphery have ON cells with highest frequency selectivity. Same color codes are used for the following figures. **C**) Covariation of time domain peak-time parameter with the respective temporal frequency selectivity for all the cells (left) and separated by ON and OFF populations (right). **D**) Covariation of time domain zero-cross parameter with the temporal frequency bandwidth for all the cells (left) and separated by ON and OFF populations (right). **E**) Covariation of time domain biphasic-index parameter with the respective temporal frequency bandwidth for all the cells (left) and separated by ON and OFF populations (right). **F**) Covariation of time domain peak-time with the temporal frequency selectivity for all the cells (left) and separated by ON and OFF populations (right). **G**) Relation between the frequency selectivity and frequency bandwidth for all the RGCs analyzed. Top and right histograms show the distribution of frequency bandwidth and frequency selectivity, respectively, for each cell population observing significant differences between them (see text). **H**) Condensed representation of **E** where each point represents a temporal cluster. Additionally, we matched properties of the frequency domain with the respective temporal profiles. Frequency selectivity is related with the cell biphasic-index. On the other side, the response latency given by peak-time and zero-cross parameters is related with the frequency bandwidth.

**Figure 6:**
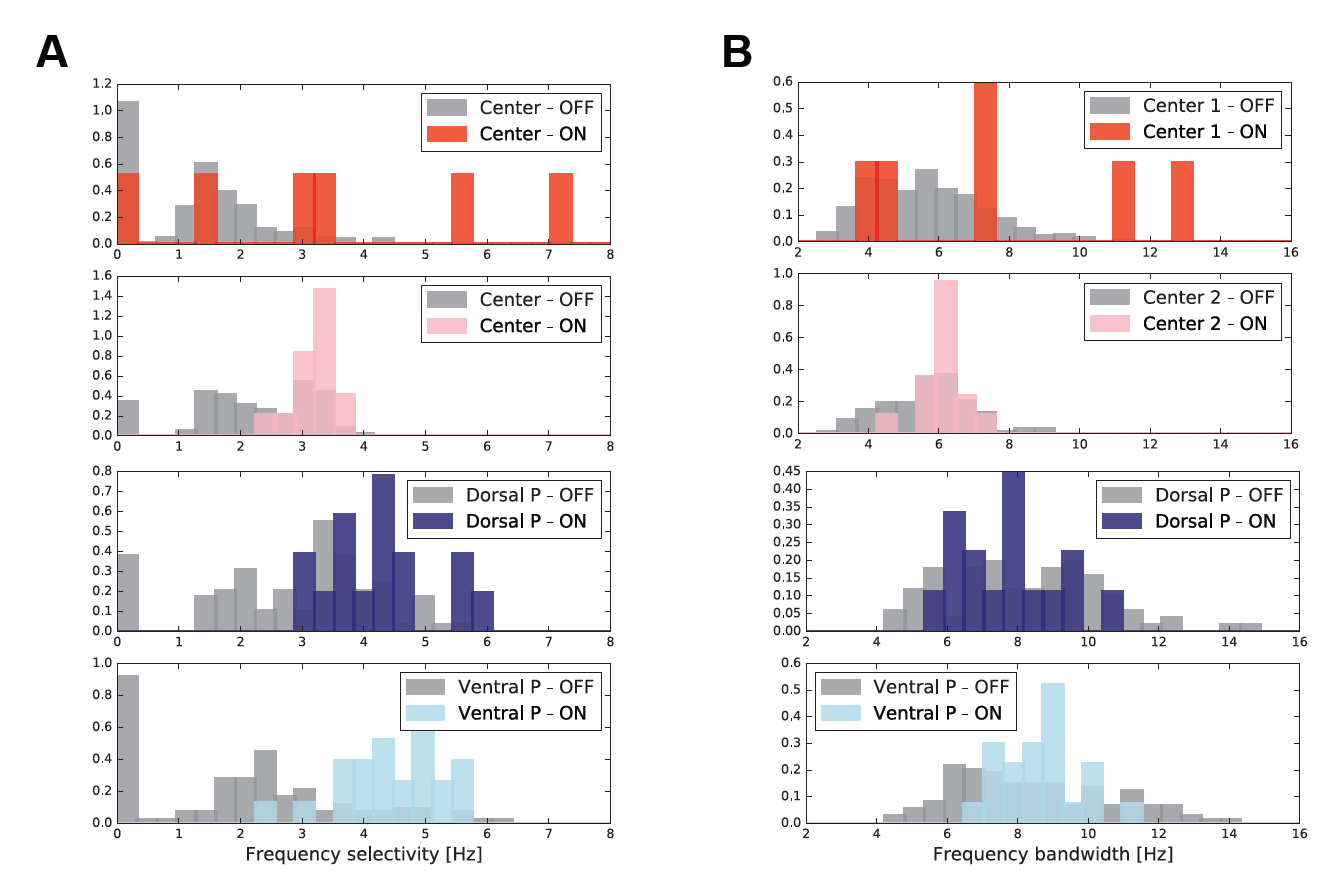
**A**) Comparison of temporal frequency selectivity between ON and OFF cell populations encountered in the same retina piece. **B**) Comparison of temporal frequency bandwidth between ON and OFF cell populations encountered in the same retina piece.

We also observed whether peak-time parameter in the time-domain is correlated with the temporal frequency selectivity (see Fig. 5**C**). No correlation is observed between these two parameters (r-value = 0.14, p-value*<*1e-3). Looking at the ON / OFF functional differences (Fig. 5**C** right) we observe that ON population contains the highest frequency selectivities with the slowest peak-time response, having in the periphery faster responses and higher frequency selectivities.

In opposite, fast and shorter temporal responses increase the temporal frequency bandwidth. A global correlation is evidenced between peak-time parameter in the time-domain with the temporal frequency bandwidth (r-value = 0.60, p-value*<*1e-5, see Fig. 5**D**). Similarly, there is also exist correlation between the zero-cross time-domain parameter with the frequency bandwidth (r-value = 0.58, p-value*<*1e-5, see Fig. 5**E**). Analyzing ON and OFF cell populations separately, both for center and periphery, Fig. 5**D**-right presented slow ON cells (larger peak-time values). ON cell population in the peripheral retina is condensed on small set of short temporal responses (low zero-cross values), by the contrary, same cell population in the central retina exhibits a larger range of temporal response durations, see e.g., Fig. 5**E**-right.

Now relating the biphasity of temporal cell response with the temporal frequency selectivity (see Fig. 5**F**), we observe a strong correlation between these two parameters (r-value = 0.68, p-value*<*1e-5). As we previously mentioned in the temporal analysis (see Fig. 4), there are significant differences between the BI of the different retina analyzed, where central retina presents a tendency to have cells less biphasic. Moreover, analyzing ON and OFF cell populations separately, we observe that ON cells (see Fig. 5**F**-right) are characterized by a very high BI parameter, together with a high temporal frequency bandwidth.

Fig. 5**G** shows the relation between frequency selectivity and frequency bandwidth for all the RGCs cells encountered in each piece of retina observing a high correlation (center: r-value = 0.73, p-value*<*1e-5; periphery: r-value = 0.67, p-value*<*1e-5). We observed that low-pass filtering have bounded frequency bandwidth for the central RGCs. By the contrary, in the periphery low-pass filtering is observed in a wide range of frequency bandwidths. These observations are accompanied by histograms placed at the top and right of the main chart. Horizontal histograms shows the distribution of the frequency bandwidth parameter for all the central and peripheral RGCs, showing different selectivities in the frequency space (ANOVA p*<*0.001). Similarly, the vertical histograms represent the distribution of frequency selectivity, which are also different for each cell population (ANOVA p*<*0.02). Analyzing the contribution of either ON or OFF cell population on this characterization (see Fig. **??G**-left), we observed that in the central retina ON cell population has the highest frequency selectivity and frequency bandwidth, which slightly differs with peripheral ON cells. In the periphery, ON cells maintain the highest frequency selectivity but they are distributed along all the frequency bandwidth.

Additionally, we wanted to observe the relation between the peak frequency selectivity and the frequency bandwidth in a condensed manner, where each point represents a temporal cluster (see Fig. 5**F**). We observe a covariation of both parameters which is more evidenced for the central regions (center: r-value = 0.81, p-value*<*0.001; periphery: r-value = 0.44, p-value*<*0.05). In this condensed representation we matched the cell profiles observed in the temporal domain with their characteristics in the frequency domain (see also Fig. 5**E**). From this analysis, and what is observed in **B**, **C** and **D**, we can conclude that large zero-cross values in the temporal domain are associated to small frequency bandwidths. Similarly, small zero-cross values are associated to large frequency bandwidths. Regarding frequency selectivity, we can observe that it is correlated with the biphasic index of the temporal response. Cells with a high biphasic index have a higher frequency selectivity compared to RGCs with a low biphasic index.

Contrasting frequency properties between ON and OFF cell populations, we observed that central retina is characterized by an ON cell population with a higher frequency bandwidth compared to OFF cells. In the periphery, distributions are similar, but ON cells exhibit a smaller variability (see Fig. **??**).

### 3.6. Linear versus non-linear response

We additionally compared the linearity of RGCs in the central and peripheral retina. To do this, we estimated the linear response convolving the checkerboard stimulus with the cell RF obtained using STA. Using he algorithm proposed by McFarland et al. (2013), we covariated each cell linear response with its firing rate as it is shown in Fig. 7**A** (red for the center and blue for the periphery). The level of nonlinearity of each cell response was obtained computing the Nonlinear-Index (NI) proposed by Nichols et al. (2013), ranging from 0 to 100, where values close to 0 indicate a high nonlinear response. The inset of graph shows the distribution of the NI obtained for all the RGCs in that retina region fitted with a Gaussian. We observed no homogeneity between the NI distribution of center regions (ANOVA, p *<* 0.001), neither peripheral regions (ANOVA, p *<* 0.001).

**Figure 7:**
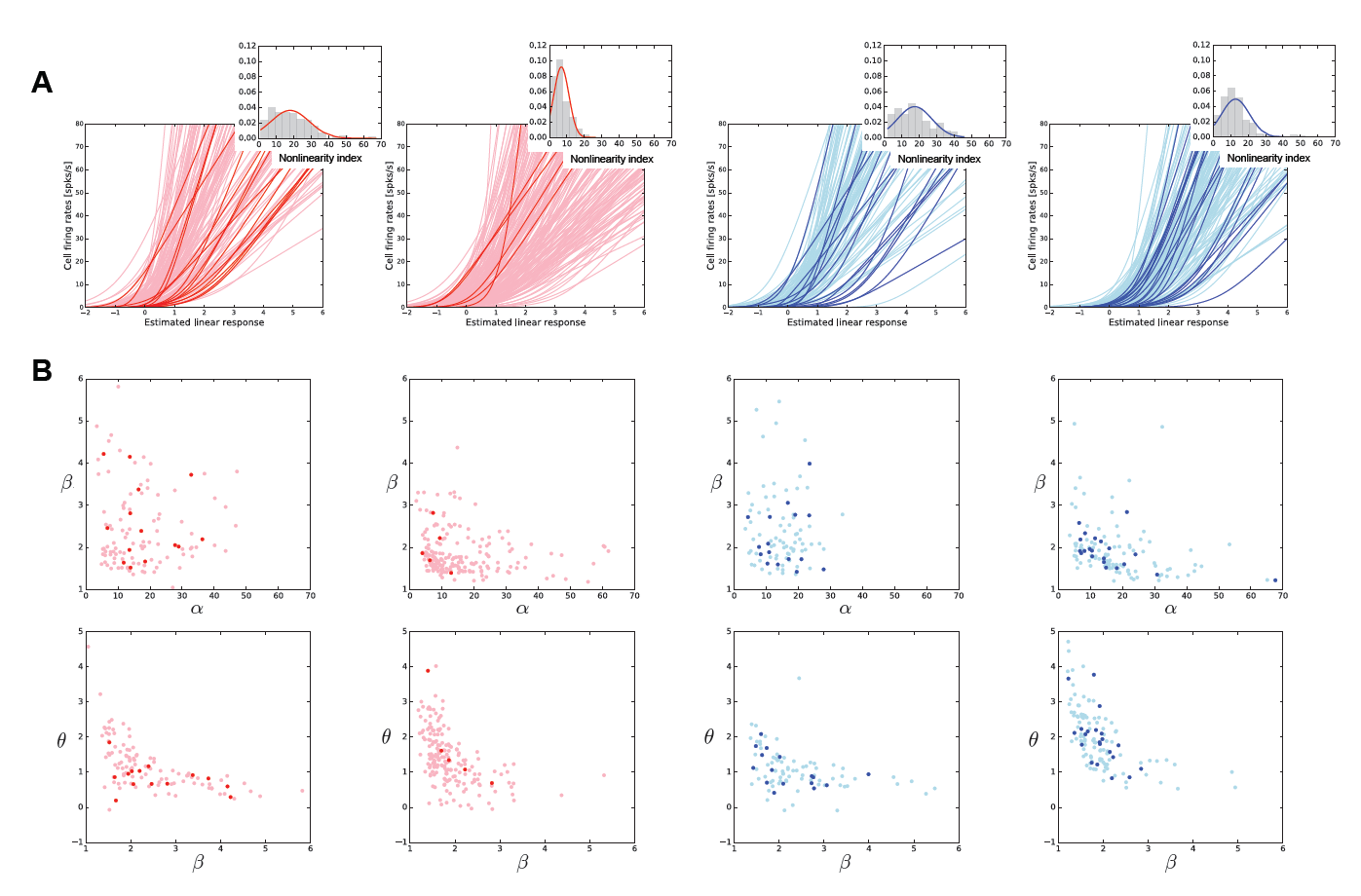
Linearity of RGC responses in the central and peripheral retina. **A**) Relation between linear cell response versus firing rate, each line represent a single cell where it is possible to evidence the nonlinear response of RGCs. Redish lines are for central regions (red: ON and light-red: OFF cell) and bluish lines for peripheral regions (blue: ON and light-blue: OFF cell). Figure insets are the distribution of NI, where values close to 0 represent a high nonlinear response. **B**) Each curve plotted in **A** was fitted by a nonlinear function with parameters *α, β* and *θ*, which values for each experiment, are here related. **A** and **B** use the same color code.

Each of the relation of cell firing rate versus linear response was fitted by the equation described in 3, and parametrized where the parameters *β* and *θ* are related to the linearity of the cell (see Fig. 7**B**). Interestingly, we observe similarities and differences between all the retina tissues, not associated to a central or peripheral category. This variety of behavior is also observed across OFF (filled colored circles) and ON (empty circles) populations.

### 3.7. Simulated retina response in a real scenario

Using the spatial and temporal profiles observed in the central and peripheral retina of the *Octodon degu*, we asked whether their functional properties have a computational and therefore ecological meaning. To do so, we simulated a central and peripheral retina selecting the linear RF of two sample temporal profile clusters per region. We considered one ON and one OFF cluster with RF sizes of 80-90um for center and 100-110um for the periphery region (OFF-ON cell respectively), which spatial and temporal profiles are shown in Fig. 8**A-B**.

**Figure 8:**
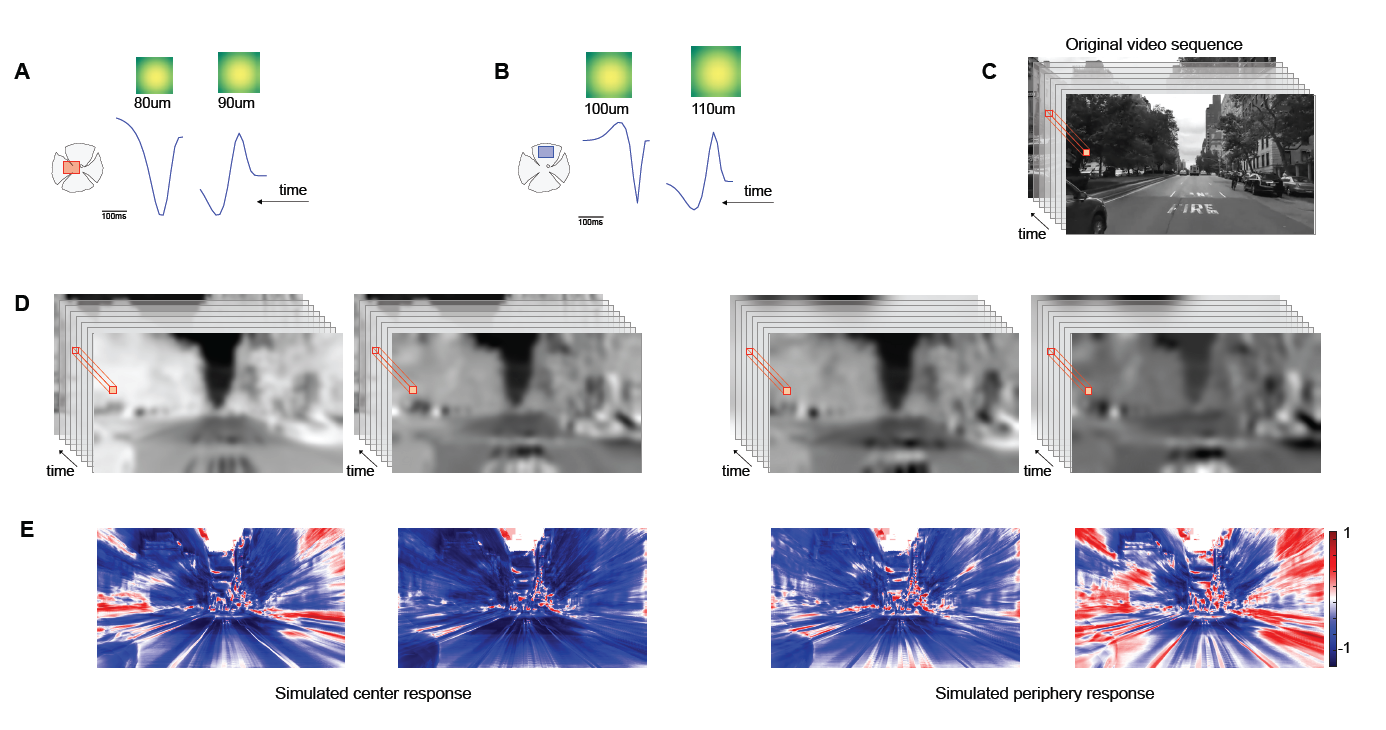
Simulated linear retina response in a real scenario. **A-B** Two sample cells were selected for the center and periphery regions. Spatial and temporal profiles are shown in the figure. **C** Input video sequence from New York city used in this simulation. **D** Each column represents the simulated RGC response for each cell shown in **A** for the center (left) and periphery (right). **E** Correlation between the input video and the simulated RGC response. Red represents a high correlated value, while blue an uncorrelated response.

An input video sequence of an approaching motion in New York city (Fig. 8**C** 640 × 480 pixels, 200 frames, 60fps) was used as input and it was convolved by two sample cells. The input video was linearly filtered by each of the RGCs shown in **A-B**, obtaining the responses shown in **C**. For each pixel of the resulting videos, we extract its temporal variation as a vector and we computed the dot product between this vector and the one obtained from the original video. This procedure allowed us to detect the spatial regions of the video mainly encoded by the clusters associated to central or peripheral RGCs. Fig. 8**E** shows the results obtained in each case, where it is possible to observe that the central retina mainly encodes the information located at the center of the image (which is mainly static), while the peripheral retina cares about the moving parts which are mostly located at the lateral borders of the original video sequence.

## 4. Discussion

Here, using a 256-MEA system, we report the functional characteristics of RGCs placed at the central and peripheral retina of a diurnal rodent (*Octodon degus*). In this animal model, it has been reported an increased cell density of photoreceptors (M and UV cones) and RGCs in a horizontal band around the optic nerve (Jacobs et al., 2003; Vega-Zuniga et al., 2013) suggesting differences in the computational strategies performed in these two regions. We analyzed the functional characteristics of central and peripheral retina patches observing important asymmetries between them, at the spatial and temporal level.

A global analysis revealed strong asymmetries between central and peripheral RGCs. In the periphery, RGCs have larger RF sizes compared to central RGCs, suggesting large integration field areas (Fig. 3). Similar results were reported in the archer fish, where RFs at the centralis stream have a resolution comparable to photoreceptors (Ben-Simon et al., 2012). Peripheral RGCs have more sustained response than the center (Fig. 2) and faster with a shorter time course, which was evidenced using both light flashes and checkerboard responses (Fig. 4). After the frequency analysis, we observed that fast and short temporal responses (smaller zero-cross and peak-time values) are related with a wider frequency bandwidth and a higher frequency selectivity, which is also a feature of peripheral RGCs (Fig. 5). These findings are fully consistent with the characteristics of higher visual areas of the visual system. For instance, Orban et al. (1986) compared central V1 and V2 visual areas versus peripheral V1 areas in the macaque monkeys reporting that receptive field sizes increase with eccentricity and the speed selectivity of V1 neurons also increases with eccentricity, which is mostly determined by the temporal frequency selectivity.

Cell characterization relies on several cell features such as morphology, physiological responses and molecular properties, which remains a challenging problem without consensus (Armañanzas and Ascoli, 2015). From a computational point of view, where the interest relies on the strategies performed by the retina, a functional characterizations based on physiological recordings seems natural. In our case, the functional type characterization was done considering the temporal profile of the linear response and functional types were obtained grouping temporal profiles following the clustering algorithm proposed by Rodriguez and Laio (2014). Alternative manners to compute functional cell types could be also consider in the analysis (Sharpee, 2013). For instance, considering aspects of information theory Sharpee (2014) proposes a methodology mainly based on the nonlinearity response of each cell. In salamander retina, Kastner and Baccus (2011) applied this methodology to find functional types, which were a result of the noise statistical properties affecting neural responses. Interestingly Kastner and Baccus (2011) found that the uniform sample of the visual scene done by different functional types, classically observed in non-human primates (Field et al., 2007; Gauthier et al., 2009), is also present in the salamander retina. Further analysis could also be done for a better functional characterization, for instance, considering motion and speed processing capabilities of each functional type (see e.g. Ben-Simon et al. (2012)).

We asked whether these asymmetries are not only observed between RGCs located at the central and peripheral retina, but also between ON and OFF populations. Regarding spatial characterization, the central pieces of retina did not show differences between RF sizes of ON and OFF cells, only the ventral periphery showed RF size of ON population larger than OFF population (Kolmogorov-Smirnof, p*<*0.02). Regarding temporal characterization, the latencies of ON cells compared to OFF cells vary along different retinal regions and the type of stimulus. Using light flashes we observed opposite behaviors in the two centers and two peripheral pieces analyzed: ON cell population can be either faster or slower than OFF cells (see Fig. 2). The results using checkerboard, and considering the peak-time parameter, revealed a faster OFF population compared to ON cells. Nevertheless, observing the zero-cross parameter related to the response duration, we observed that ON cells have shorter temporal courses compared to OFF cells in almost all the tissues analyzed (see Fig. 4). Functional differences between ON and OFF cell kinetics were previously reported by Chichilnisky and Kalmar (2002), who described in non-human primates ON cells with a faster response kinetics compared to OFF cells and ON cells with RF sizes 20% larger than OFF cells. Similarly, Zaghloul et al. (2003) sought for the origins of this asymmetry finding significant differences in the synaptic inputs between ON and OFF populations (guinea pig).

It has been also reported asymmetries between ON and OFF RGCs population in terms of transient or sustained response. Jiang et al. (2015), analyzing retinal data from non-human primates, encountered OFF neurons having higher spontaneous activities, higher peak response amplitudes and with a more sustained temporal response compared to ON neurons. Nevertheless, we only observed asymmetries between central and peripheral RGCs, not particularly associated to ON or OFF populations.

The asymmetries observed between temporal responses of ON and OFF populations can be studied in the temporal frequency domain. In general, we observed than ON population has higher frequency selectivity and higher bandwidth compared to OFF cells (see Fig. 5). We observed several RGCs acting as temporal low-pass filters, but this property is only present in OFF cells, not observed in ON cells. Considering that speed can be computed in terms of temporal and spatial frequency selectivities, we can conclude that greater RF sizes and higher temporal frequency selectivities (ON properties) exhibit a selectivity to higher speeds compared to cells with lower temporal frequency selectivities (OFF cells). Nevertheless, it seems that ON population presents a selective deficit in carrying information about moving stimuli, which deficit is not observed in OFF population (see Nichols et al. (2013)). This finding is opposite to Leonhardt et al. (2016), who reported in drosophila, that ON cells respond maximally to slower velocities compared to OFF cells suggesting that speed information is encoded by ON and OFF populations in a combined manner: at slow velocities the coding is driven by ON cells, while at high velocities by OFF cells. These results could be probably combined with our findings in terms of the temporal or spatial frequency bandwidth, where ON and OFF populations presents different distributions.

Considering the entire ON and OFF populations found in the retina pieces analyzed, we observed no significant differences between them in terms of linear / no-linear responses. Nevertheless, it has been reported in the literature that the linearity of their responses differs. For instance, Chichilnisky and Kalmar (2002) showed ON cells with nearly linear light responses and OFF cells acting more like rectifiers. Similarly, looking for the underlying mechanism to justify this behavior, Liang and Freed (2010) reported that input currents in OFF cells were more rectified than those in ON cells. This asymmetry may be an adaptation to natural scenes, that have more contrast levels below the mean than above. The differences between our findings and those encountered in the literature can be attributed mainly to the type of ON cell observed, and the differences associated to the animal model in study.

Different functionalities of the central and peripheral vision are attributed to ecological needs involved in behavior and survival. Central vision encodes fine details of the visual scene, while peripheral vision is related to coarse spatial and temporal frequency information mostly related to generate alert messages needed for survival. Behavioral studies in humans related with object recognition revealed a better performance when the object is placed within the 1-2^*?*^ of the visual field (Musel et al., 2013), similar is the case for face recognition (Arcaro et al., 2009). Nevertheless, studies in fMRI evidence that humans confronted to natural scenes their visual peripheral regions are more activated suggesting a combined processing strategies for the understanding of the visual world (Nasr et al., 2011; Loschky et al., 2017). Interestingly, using a deep neural network the authors in Wang and Cottrell (2017) have shown than information coming from peripheral vision is as crucial as central vision for scene recognition.

This consequence in behavior and recognition strategies of our visual system, initially obtained from the big variety of functional RGC types located in the retina, that according to our study, even within the same animal specie, must be done carefully selecting the same retina area of analysis. Predictive coding theory hypothesize that the role of neural encoding is to remove predictable information from the environment and only leave what cannot be predicted. The work of Nirenberg et al. (2010) goes in this direction suggesting that heterogeinity in RGC types is needed to predictive coding in different types of environments. Moreover, Gjorgjieva et al. (2017) recently reported that optimal coding in the retina is reached not by ON or OFF isolated neuron population, by a mixture of ON / OFF neurons suggesting a combined coding strategy.

## 5. Acknowledgments

Financial support: FONDECYT 1140403 and 1150638, CONICYT-Basal Project FB0008; Grant ICM-P09-022-F supported by the Millenium Scientific Initiative of the Ministerio de Economia, Desarrollo y Turismo (Chile); ECOS-Conicyt C13E06; ONR Research Grant # N62909-14-1-N121. We would also like to thank to Dr. Ronny Vallejos and Dr. Felipe Osorio for their valuables comments about the statistical analysis of the results.

